# Antimicrobial efficacy of neem and liquorice with chlorhexidine on Streptococcus sanguis, Streptococcus mutans, Lactobacillus and Actinomyces naeslundii – An In Vitro Study

**DOI:** 10.1101/2020.09.23.311019

**Authors:** Saad M Alqahtani

## Abstract

The study was to formulate 2% neem and 2% liquorice mouthwashes and to compare the antimicrobial efficacy of these mouthwashes with the standard 0.2% chlorhexidine mouthwash. Alcoholic solution was prepared and added to neem mixture and liquorice mixture separately and made up to a volume of 16000 ml with purified water. Nine dilutions of each drug were done with Brain heart infusion broth (BHI) for MIC. Culture suspension was added in each serially diluted tube of 200 μl. The tubes were incubated for 24 hours and observed for turbidity. Minimum inhibitory concentration (MIC) of 2% neem, 2% liquorice and 0.2% chlorhexidine against Lactobacillus, Actinomyces naeslundii, Streptococcus sanguis, Streptococcus mutans is determined by serial dilution analysis. Streptococcus mutans shows sensitivity to all three mouthwashes at a concentration starting from 0.2 μg/ml. Lactobacillus shows sensitivity to neem and chlorhexidine mouthwashes at a concentration starting from 1.6 μg/ml, whereas liquorice is effective at a concentration starting from 3.125 μg/ml. Streptococcus sanguis shows sensitivity to chlorhexidine and liquorice mouthwashes at a concentration starting from 25 μg/ml, whereas it shows sensitivity to neem at a concentration starting from 50 μg/ml. Actinomyces naeslundii shows sensitivity to chlorhexidine and neem mouthwashes at a concentration starting from 1.6 μg/ml, whereas it shows sensitivity to liquorice at a concentration starting from 3.125μg/ml. Analysis showed an inhibition of all the four strains by the mouthwashes. The MIC for the studied mouthwashes was found to be similar to that of 0.2% chlorhexidine.

## INTRODUCTION

Plaque is an organized mass, consisting mainly of microorganisms, that adheres to teeth, prostheses, and oral surfaces and is found in the gingival crevice and periodontal pockets. It also comprises of an organic compound consisting of bacterial by-products such as enzymes, materia alba, desquamated cells, and inorganic components such as calcium and phosphate [1]. Gingivitis, which has a direct association with dental plaque [2], affects the oral health of 70%–100% of the population across the world [3-5]. Plaque-induced gingivitis is an inflammatory response of the gingival tissues resulting from bacterial plaque accumulation located at and below the gingival margin [6]. It does not result in clinical attachment loss or directly cause tooth loss [7]. It is the most common form of gingival disease. Patients may notice symptoms that include bleeding with tooth brushing, blood in saliva, gingival swelling and redness, and halitosis in the case of established forms [8]. Experimental studies done on plaque induced gingivitis have shown the first empiric evidence that accumulation of microbial biofilm on clean tooth surfaces reproducibly induces an inflammatory response in the associated gingival tissues and removal of plaque leads to disappearance of the clinical signs of inflammation [9]. Plaque control is the regular removal of microbial plaque and the prevention of its accumulation on the teeth and adjacent gingival surfaces. The utilization of antimicrobial mouth rinses has been considered a useful adjunct to mechanical plaque control [10]. Tooth brushing is the most popular self-performed oral hygiene method to mechanically remove dental plaque. However, this mechanical approach by most individuals is often not sufficiently effective [11]. Chlorhexidine (0.2%) is considered the gold standard chemical plaque control agent because of its clinical efficacy. It is one of the most investigated compounds in dentistry and has been proven to inhibit plaque regrowth and the development of gingivitis by the first definitive experimental study [12].

Herbal medicines, derived from botanical sources, have been applied in dentistry for a long history to inhibit microorganisms, reduce inflammation, soothe irritation, and relieve pain [13]. Literature reports that a considerable number of herbal mouthwashes have achieved encouraging results in plaque and gingivitis control. When compared with antimicrobial activity of synthetic commercial mouthwashes, herbal mouthwashes can exhibit additional anti-inflammatory and antioxidant properties, which could further improve periodontal health [14]. Neem leaf and its constituents have been identified to exhibit immunomodulatory, antiinflammatory, antihyperglycaemic, antiulcer, antimalarial, antifungal, antibacterial, antiviral, antioxidant, antimutagenic and anticarcinogenic properties. The antibacterial effect of neem mouthwash against salivary levels of Streptococcus mutans and Lactobacillus has been demonstrated [15]. Liquorice extracts and its principal component, glycyrrhizin exhibits useful properties such as anti-inflammatory, antiviral, antimicrobial, antioxidative, anticancer, immunomodulatory, hepatoprotective and cardioprotective effects [16]. Licorice mouthwash has shown its effectiveness in reducing plaque accumulation and gingival inflammation along with no discoloration of teeth or unpleasant taste [17].

The present study was undertaken to formulate 2% neem and 2% liquorice mouthwashes and seeks to compare the antimicrobial efficacy of these mouthwashes with the standard 0.2% chlorhexidine mouthwash.

## MATERIALS AND METHODS

The neem leaf extract and liquorice extract were obtained from Amsar Private Limited, Bardez, Goa, India, and the preparation of mouthwash was done at AVN Arogya Ayurvedic Hospital, Madurai, India. (Fig.1)

**Fig. 1:**
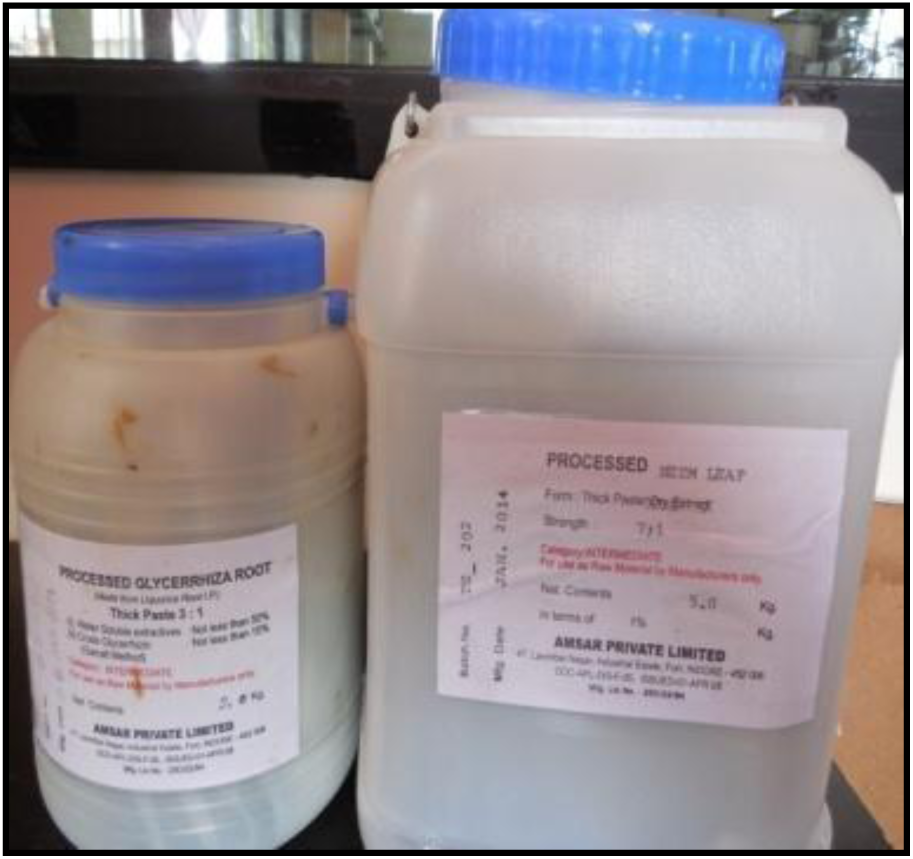
Neem and Liquorice extracts.

All the materials were dispensed as per the list using suitable balances. Neem extract was dissolved in water separately. Sodium benzoate and benzoic acid were dissolved in alcohol (96%). Saccharin sodium, sorbitol and glycerin were dissolved in water and then transferred to the alcoholic solution. The above alcoholic solution was added to the neem mixture and made up to a volume of 16000 ml with purified water (Fig.2) to the filtered solution (Fig.3), peppermint oil was added and stirred.

**Fig. 2:**
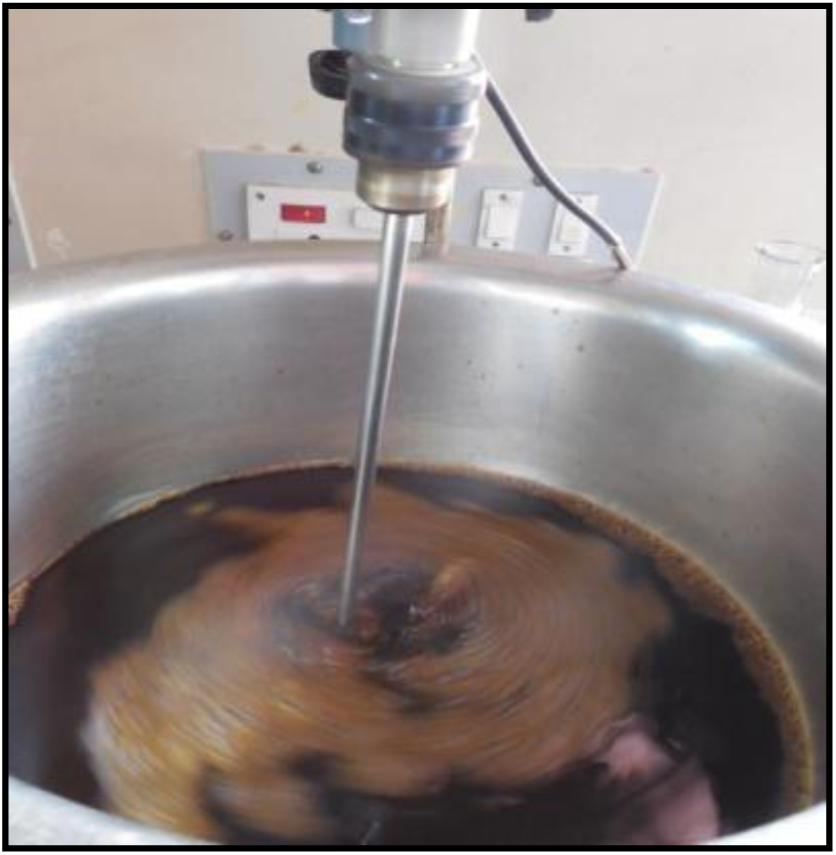
Mixing of ingredients with stirrer.

**Fig. 3:**
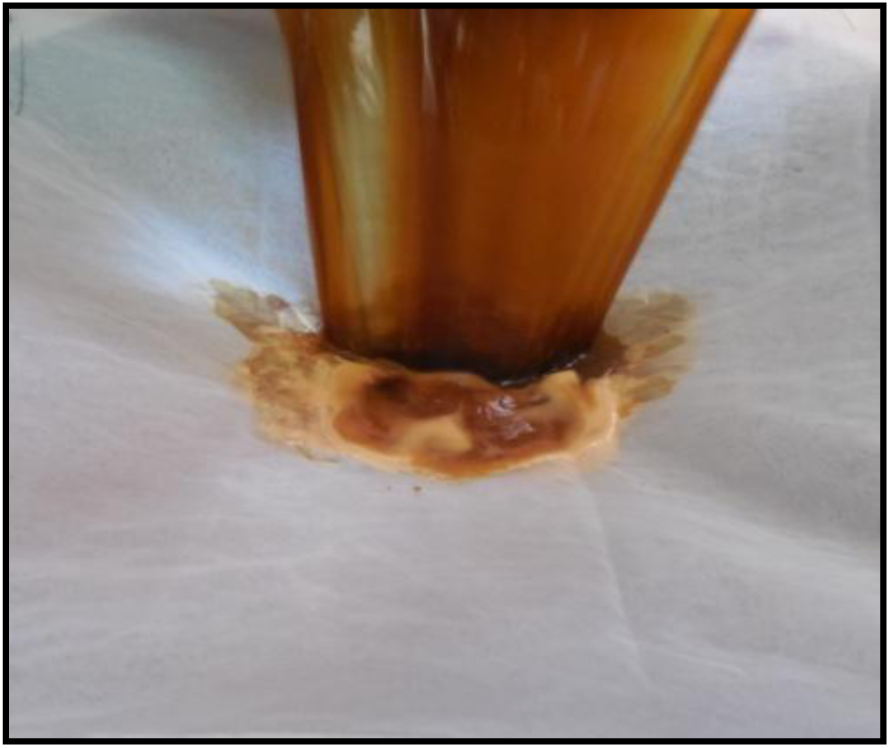
Filtering the prepared mouthwash.

All the materials were dispensed as per the list using suitable balances. Liquorice extract was dissolved in water separately. Sodium benzoate, citric acid, saccharin sodium, sorbitol and glycerin were dissolved in water and then transferred to the alcoholic solution. The above alcoholic solution was added to the liquorice mixture and made up to a volume of 16000 ml with purified water. To the filtered solution, peppermint oil was added and stirred. Commercially available 0.2% chlorhexidine (Hexifresh mouthwash) was used. Both the prepared mouthwashes were stored in seaparate dark opaque bottles. (Fig.4) Minimum inhibitory concentration (MIC) of 2% neem, 2% liquorice and 0.2% chlorhexidine against Lactobacillus, Actinomyces naeslundii, Streptococcus sanguis, Streptococcus mutans is determined by serial dilution analysis.

**Fig. 4:**
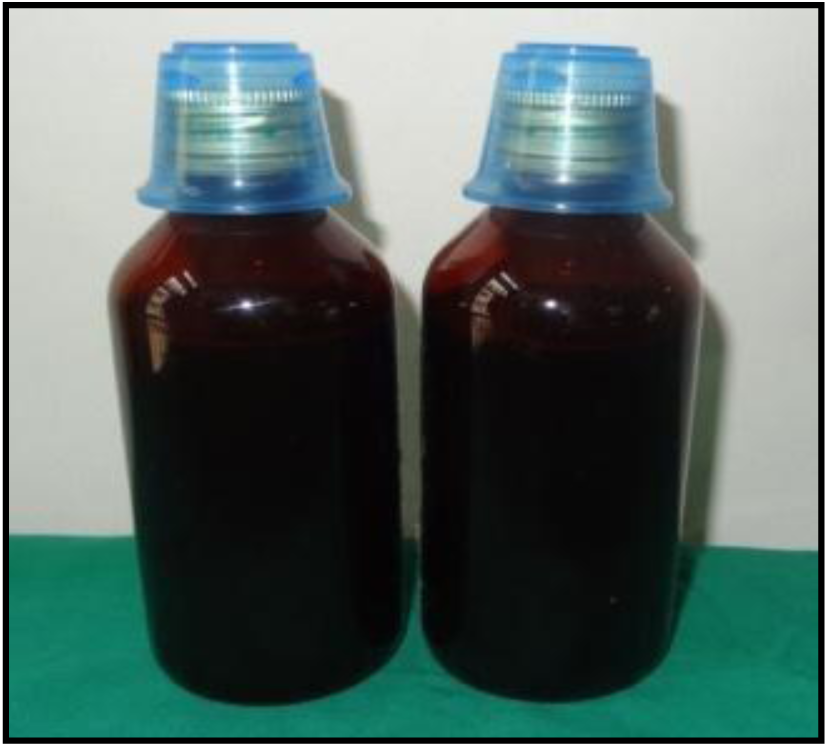
Neem and liquorice mouthwashes in opaque bottles.

### MIC procedure

Nine dilutions of each drug were done with Brain heart infusion broth (BHI) for MIC (Fig.5) In the initial tube 20 μl of drug was added into 380microliter of BHI broth. For further dilutions 200μl amount of BHI broth was dispensed into the next 9 tubes separately. Then from the initial tube 200 μl was transferred to the first tube containing 200 μl of BHI broth. This was considered as 10^−1^ dilution. From 10^−1^ diluted tube 200 μl was transferred to second tube to make 10^−2^ dilution. The serial dilution (Fig.6) was repeated up to 10^−9^ dilution for each drug. Out of the stock cultures that were maintained, 5 μl was taken and added into 2 ml of BHI (brain heart infusion) broth. Above culture suspension was added in each serially diluted tube of 200 μl. The tubes were incubated for 24 hours and observed for turbidity.

**Fig. 5:**
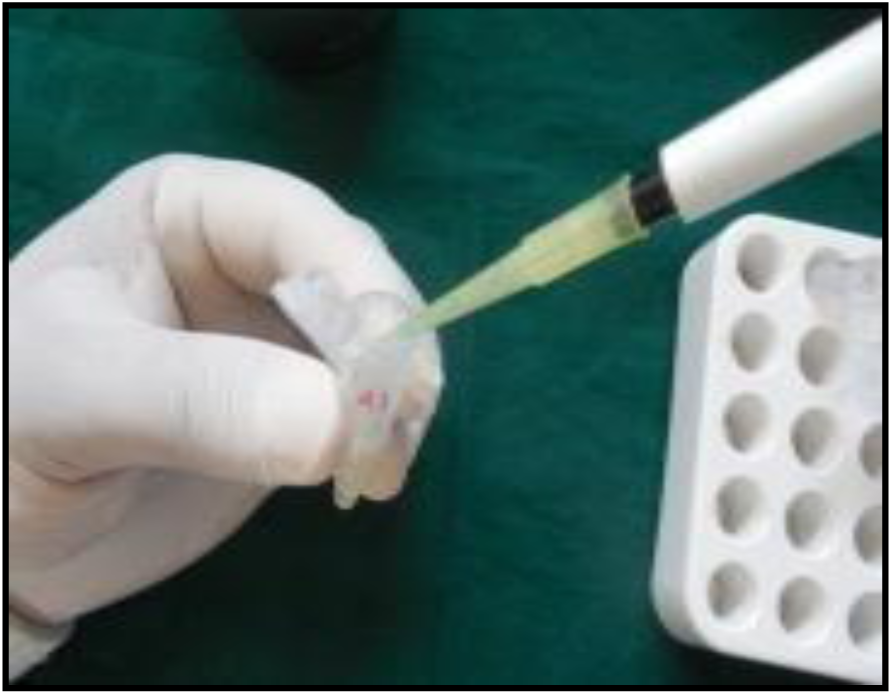
Dispensing BHI broth into tubes.

**Fig. 6:**
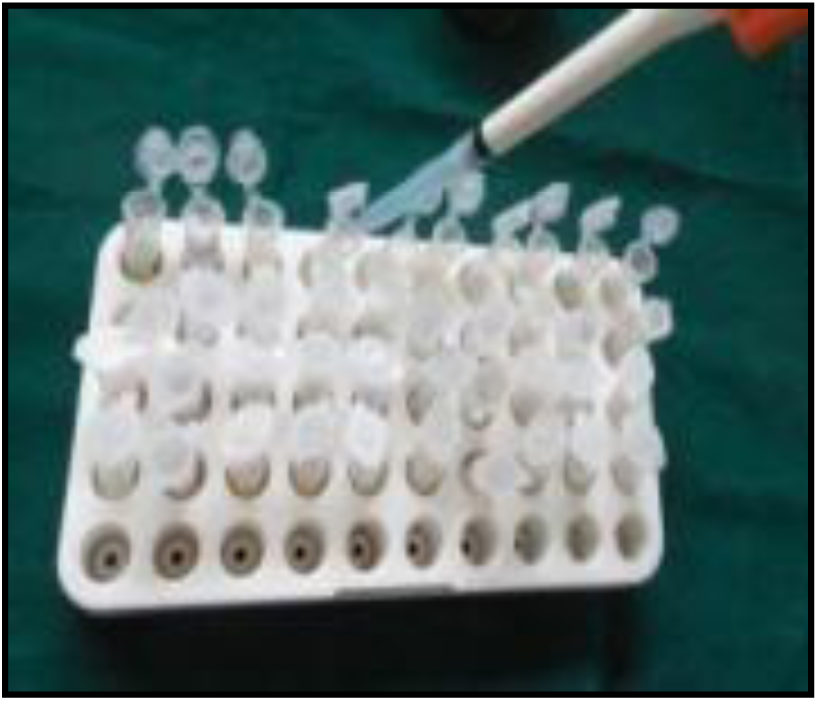
Serial dilution of the drug.

## RESULTS

In the present study, we formulated 2% neem, 2% liquorice and a commercially available 0.2% chlorhexidine mouthwash. An in vitro analysis of the antibacterial efficacy of the mouthwashes was performed, followed by an in vivo analysis of the clinical efficacy in 90 subjects randomized into three groups. During the study, five participants did not complete the study and were excluded from the analysis (2 from neem, 2 from liquorice and 1 from chlorhexidine mouthwash groups). Table 1 shows the in vitro analysis of the antimicrobial sensitivity test of Streptococcus mutans, Streptococcus sanguis, Actinomyces naeslundii and Lactobacillus to neem, liquorice and chlorhexidine mouthwashes by serial dilution analysis.

**Table: 1:** In vitro analysis:

| Streptococcus mutans |  |  |  |  |  |  |  |  |  |  |  |
| --- | --- | --- | --- | --- | --- | --- | --- | --- | --- | --- | --- |
| S.N | Mouthwash | 100<br>µg/ml | 50<br>µg/ml | 25<br>µg/ml | 12.5<br>µg/ml | 6.25<br>µg/ml | 3.125<br>µg/ml | 1.6<br>µg/ml | 0.8<br>µg/ml | 0.4<br>µg/ml | 0.2<br>µg/ml |
| 1. | Neem | S | S | S | S | S | S | S | S | S | S |
| 2. | Liquorice | S | S | S | S | S | S | S | S | S | S |
| 3. | Chlorhexidine | S | S | S | S | S | S | S | S | S | S |
| Lactobacillus |  |  |  |  |  |  |  |  |  |  |  |
| 1. | Neem | S | S | S | S | S | S | S | R | R | R |
| 2. | Liquorice | S | S | S | S | S | S | R | R | R | R |
| 3. | Chlorhexidine | S | S | S | S | S | S | S | R | R | R |
| Streptococcus sanguis |  |  |  |  |  |  |  |  |  |  |  |
| 1. | Neem | S | S | R | R | R | R | R | R | R | R |
| 2. | Liquorice | S | S | S | R | R | R | R | R | R | R |
| 3. | Chlorhexidine | S | S | S | R | R | R | R | R | R | R |
| Actinomyces naeslundii |  |  |  |  |  |  |  |  |  |  |  |
| 1. | Neem | S | S | S | S | S | S | S | R | R | R |
| 2. | Liquorice | S | S | S | S | S | S | R | R | R | R |
| 3. | Chlorhexidine | S | S | S | S | S | S | S | R | R | R |

Streptococcus mutans shows sensitivity to all three mouthwashes at a concentration starting from 0.2 μg/ml. Lactobacillus shows sensitivity to neem and chlorhexidine mouthwashes at a concentration starting from 1.6 μg/ml, whereas liquorice is effective at a concentration starting from 3.125 μg/ml. Streptococcus sanguis shows sensitivity to chlorhexidine and liquorice mouthwashes at a concentration starting from 25 μg/ml, whereas it shows sensitivity to neem at a concentration starting from 50 μg/ml. Actinomyces naeslundii shows sensitivity to chlorhexidine and neem mouthwashes at a concentration starting from 1.6 μg/ml, whereas it shows sensitivity to liquorice at a concentration starting from 3.125μg/ml.

## DISCUSSION

The formation of dental plaque, the primary etiological factor for periodontal diseases is characterized by the initial adherence of limited number of pathogenic bacteria to the salivary pellicle and progressive accumulation and growth of complex flora. There is a direct relationship between the presence of dental plaque and the development of gingivitis, which is characterized by the inflammation of gingiva without clinical attachment loss [10]. Plaque control usually means preventive measures aimed at removing dental plaque and preventing it from recurring. Complete removal of bacterial plaque from the dento-gingival region is the most effective method of preventing gingivitis and periodontitis and the effective removal of dental plaque is important for dental and periodontal health. This can be accomplished through mechanical and chemical measures [18].

Tooth brushing is the most accepted oral hygiene practice. Chemical inhibitors of plaque and calculus incorporated in mouthwashes or dentifrices also play an important role in plaque control as adjuncts to mechanical cleansing [18]. The side effects associated with the commercially available plaque control agents has ushered an era of herbal alternatives. Neem and Liquorice are traditional herbs widely used in India and South Asia from time immemorial for maintaining healthy teeth and gingiva.

Neem is rich in a vast array of biologically active compounds that are chemically distinct and structurally multifaceted. Neem twigs are used for brushing and the leaves of neem have been effectively used in the treatment of gingivitis and periodontitis [19]. Licorice or liquorice (Glycyrrhizaglabra), is a perennial herb which possesses sweet taste and has extensive pharmacological effects [20]. Being indigenous, cheap and readily available, neem and liquorice can be definitely expected to have better patient compliance and acceptability. Due to the aforementioned potential benefits, we aimed at formulating 2% neem and 2% liquorice mouth rinses. Their efficacy in vitro was compared with a commercially available 0.2% chlorhexidine mouthwash. We assessed the antimicrobial efficacy of neem, liquorice and chlorhexidine *in vitro* using the serial dilution method with pre-cultured *Streptococcus mutans, Streptococcus sanguis, Actinomycesnaeslundii* and *Lactobacillus*, which are non-specific opportunistic pathogens that can induce gingivitis. Neem exhibited potent antimicrobial effect against Streptococcus mutans, Streptococcus sanguis, Actinomycesnaeslundii and Lactobacillus at a concentration of 0.2, 50, 1.6 and 1.6 μg /ml respectively.

Nayak A *et al* [21] observed inhibition of *E. faecalis, S. mutans, C*.*albicans* by alcoholic neem extract at 1.88%, 7.5% and 3.75% respectively and the aqueous neem extract at 7.5%. Maragathavalli S. *et al* [22] demonstrated inhibition of *Bacillus pumillus, Pseudomonas aeruginosa* and *Staphylococcus aureus* by the methanol and ethanol extracts of neem. Widowati *et al* [23] in their study concluded that the neem stick extract had a higher antibacterial effect on *Streptococcus mutans* than the neem leaf extract. Chloroform extracts of neem were identified to inhibit *Streptococcus mutans, Streptococcus salivarious* and *Fusobacteriumnucleatum* by Packialakshmi *et al* [15] The minimum inhibitory concentration of acetonic extract of neem for *Streptococcus sobrinus* was observed to be 0.05% (w/v) by M Bhuiyan *et al* [24] In a study by Prashant *et al* [25] with neem extract, maximum zone of inhibition on *Streptococcus mutans* was observed at 50% concentration with minimal effect on *Streptococcus mitis, Streptococcus salivarius* and *Streptococcus sanguis*.

Our study revealed Liquorice to be a potent inhibitor of *Streptococcus mutans, Streptococcus sanguis, Actinomycesnaeslundii* and *Lactobacillus* at a concentration of 0.2, 25, 3.125 and 3.125 μg /ml respectively.

Earlier studies on the antimicrobial effect of liquorice by Manoj. M. Nitalikar *et al* [26] with gram positive (*Bacillus subtili* and *Staphylococcus aureus*) and gram negative (*E*.*coli* and *Pseudomonas aeruginosa*) bacteria and M.H. Shirazi *et al* [27] with *Salmonella typhi, Salmonella paratyphiB, Shigellasonni, Shigellaflexneri* and *Enterotoxigenic*. *E. coli* and Vivek K. Gupta *et al* [28] with *Antimycobacterium* have proven the inhibitory effect of liquorice on microorganisms. In our study *Streptococcus mutans, Streptococcus sanguis, Actinomycesnaeslundii* and *Lactobacillus* were inhibited by chlorhexidine at concentration of 0.2, 25, 1.6 and 1.6 μg /ml respectively. This is in accordance with the antimicrobial efficacy of chlorhexidine previously reported by W.W. Briner *et al* [29] on *Streptococci* and *Actinomyces* and W.G Wade *et al* [30] on 355 subgingivalmicro organism isolates. Studies reported by T.D. Hennessely *et al* [31] on gram positive cocci and Sigrun Eick *et al* [32] on *Streptococci, Enterobacteria, Candida albicans, Porphyromonasgingivalis, Aggregatibacteractinomycetemcomitans*, and *Fusobacteriumnucleatum* have reported antimicrobial efficacy of chlorhexidine at a concentration of 0.19 to 2.0μg /ml and 0.01% to 0.50% respectively.

## CONCLUSION

1. Ruminating on the substantial therapeutic benefits of neem and liquorice, we prepared 2% neem and 2% liquorice mouthwashes and compared with 0.2% chlorhexidine mouthwash.
2. An in vitro analysis of the formulated mouthwashes, to test the effect on primary colonizers like Streptococcus mutans, Streptococcus sanguis, Lactobacillus and Actinomyces naeslundii showed an inhibition of all the four strains by themouthwashes.
3. The MIC for the studied mouthwashes was found to be similar to that of 0.2% chlorhexidine.

